# Natural monobacterial environments modulate viral infection in *Caenorhabditis elegans*

**DOI:** 10.1101/2023.06.13.544807

**Authors:** Rubén González, Marie-Anne Félix

## Abstract

The microbes associated with an organism play a pivotal role in modulating their host susceptibility to viral infections. However, the influence of individual microbes on viral infections is not well understood. Here, we examined the impact of 67 naturally bacterial associates on *Caenorhabditis elegans* susceptibility to Orsay virus. Our findings reveal that viral infection of *C. elegans* is significantly influenced by monobacterial environments. The majority of tested natural bacterial environments reduced *C. elegan*s viral infection while some increased susceptibility compared to an *Escherichia coli* environmental reference. The reduction in viral infection is not caused by degradation of the virions or poor nutrition of *C. elegans* by the bacteria. The reduction in viral infection does not require known antiviral responses, including RNA interference and transcriptional regulation downstream of the RIG-I homolog DRH-1. However, one bacterium, *Lelliottia* JUb276, reduced susceptibility but does not induce resistance to viral infection in *drh-1* mutants. Our research underscores the importance of considering the natural biotic environment in studies of viral infections and provides insights for future research on host-microbes-virus interactions and viral immunity.

**AUTHOR CONTRIBUTIONS:** Conceptualization: RG, MAF; Formal analysis: RG; Investigation: RG; Resources: MAF; Data Curation: RG; Writing - Original Draft: RG, MAF; Writing - Review & Editing: RG, MAF; Visualization: RG; Supervision: MAF; Project administration: RG, MAF; Funding acquisition: RG, MAF.

## INTRODUCTION

The biotic environment of an organism plays a key role in shaping various of its traits, including immune responses (1). In particular, microbes can influence host interactions with pathogenic viruses (2). In some cases, microbes can enhance their host immunity and reduce viral replication and infectivity, thus providing protection against infection (3, 4, 5, 6). However, conversely, some microbes can increase their host susceptibility to viral infections, some of them might even be crucial for new viruses to successfully infect their hosts (7, 8, 9). In addition to the intricate interactions between the host and its commensal and pathogenic microbes, there are complex relationships among the individual microbes that form the associated microbial community. The study of simplified pathosystems and associated microbes would facilitate the understanding of these interactions and the identification of mechanisms by which hosts’ immunity to viral infections is regulated. The nematode *Caenorhabditis elegans*, widely used as a model organism, presents an ideal model system to study microbe–host interactions, enabling insightful discoveries over the past 20 years (10). One key feature of *C. elegans* for such studies is that it is possible to free the animals of associated microbes using a bleaching treatment to which worm embryos resist, and then reassociate them at will with one or several microbes.

The natural habitat of *C. elegans* is rotting fruit or other decomposing vegetal matter, which is rich in bacteria. These bacteria not only serve as a primary food source for the nematode but also constitute part of its biotic environment (11, 12). In the last decade, bacteria associated with *C. elegans* in its natural habitats have been characterized, revealing their presence in the nematode’s external environment, intestinal lumen, and as a fraction that is lysed and digested as food (13, 14, 15). Bacterial environments have been shown to affect the nematode’s metabolism, signaling pathways, and immune responses (16, 17, 18, 19). While associated bacteria can protect *C. elegans* from extracellular pathogens like bacteria or fungi (20, 21, 22, 23, 24, 25), their effect in modulating interactions with intracellular pathogens, including viruses, remains unexplored.

The only known naturally occurring virus in *C. elegans* is the Orsay virus (OrV) (26). OrV is a positive single-stranded RNA virus with a bipartite genome similar to that of Nodaviruses. This virus enters new hosts when the nematode ingests the virions while feeding and, once inside the host intestinal lumen, it infects the intestinal cells. Infected intestinal cells produce more virions that are released and transmitted horizontally to other individuals. OrV infection affects the nematode’s fitness by reducing and delaying reproduction (27). *C. elegans* lacks immune components found in other animals, but is able to mount an immune response against viral infections through a range of mechanisms (28, 29). These known mechanisms include the use of RNA interference (RNAi) and uridylation responses to target viral RNA for degradation, ubiquitin-mediated pathways that target viral proteins for degradation, and transcriptional responses to activate specific genes in response to infection. This transcriptional response includes a specific immune response to *C. elegans*’s intestinal intracellular pathogens (i.e., microsporidia or viruses) known as the Intracellular Pathogen Response (IPR) (30). The IPR involves the upregulation of a limited number of genes, including genes belonging to the *pals* gene family (whose biochemical function is currently unknown) and genes predicted to encode ubiquitin ligase components. The IPR is dependent on the helicase DRH- 1, a RIG-I family member, which is likely a viral sensor and is also essential for the RNAi response. Downstream of DRH-1, the IPR is partially regulated by the transcription factor ZIP-1 (31, 32, 33). Severe infection triggers a general ‘biotic stress response’, which involves the upregulation of stress response genes (such as *lys-3*) (34). Many mechanisms and components involved in *C. elegans*’s immune response to Orsay virus are evolutionary conserved (31, 32, 34, 35). Therefore, studying these responses can provide valuable insights into virus-host interactions in other organisms.

In this study, we investigated the effect of bacterial environments on *C. elegans* susceptibility to OrV. We focused on monocultures of bacterial clones isolated from *C. elegans* natural habitats (13, 14, 15, 36, 37). Our objective was to determine whether the bacterial environment affects the nematode’s response to the virus and gain a deeper understanding of how specific bacterial strains may modulate host susceptibility to viruses. By investigating the interplay between *C. elegans*, its natural bacterial environment, and the Orsay virus, our study aims to provide novel insights into the role of the microbial context in modulating host-pathogen interactions and reveal the potential for discovering previously unknown defense responses that combat viral infections.

## RESULTS

### Single bacterial environments modulate *C. elegans* susceptibility to viral infection

We examined the impact of 67 natural bacterial strains (Supplementary Table 1) on the susceptibility of *C. elegans* to OrV. For this initial screen, we used animals carrying the *pals-5::GFP* reporter of intracellular infection (38). We conducted the screen in three experimental blocks (Supplementary Figure 1), testing three populations of ≈ 100 animals per bacterial environment and including *Escherichia coli* OP50 in each block as the reference. We scored the proportion of animals activating the *pals-5* reporter with and without viral exposure. None of the bacteria triggered *pals-5* reporter activation in the absence of virus. Based on the relative *pals-5* activation values compared to the *E. coli* OP50 reference (Figure 1, upper panel), we categorized the bacterial strains in three groups: 54 repressors of *pals-5* reporter activation (significant lower activation than OP50), 5 enhancers (significant higher activation than OP50), and 8 bacteria with an effect similar to that of the *E. coli* environment. A higher number of bacteria were thus repressive compared to *E. coli* than expected by chance (χ2 = 67.552, df = 2, *P* < 0.001). We tested some strains across different blocks, verifying that their effect on viral infections was reproducible (Supplementary Figure 2A); all bacteria with a relative rate higher than one were examined again within the same block, successfully replicating the observations of the initial screen (Supplementary Figure 2B). We wondered whether the bacterial environments prevented *pals-5* reporter activation under other types of stressing conditions, such as heat stress (30). The *pals-5* reporter could be activated by heat stress in these bacterial environments, showing that the suppression was specific of the response to viral infection (Supplementary Table 1).

**Figure 1.**
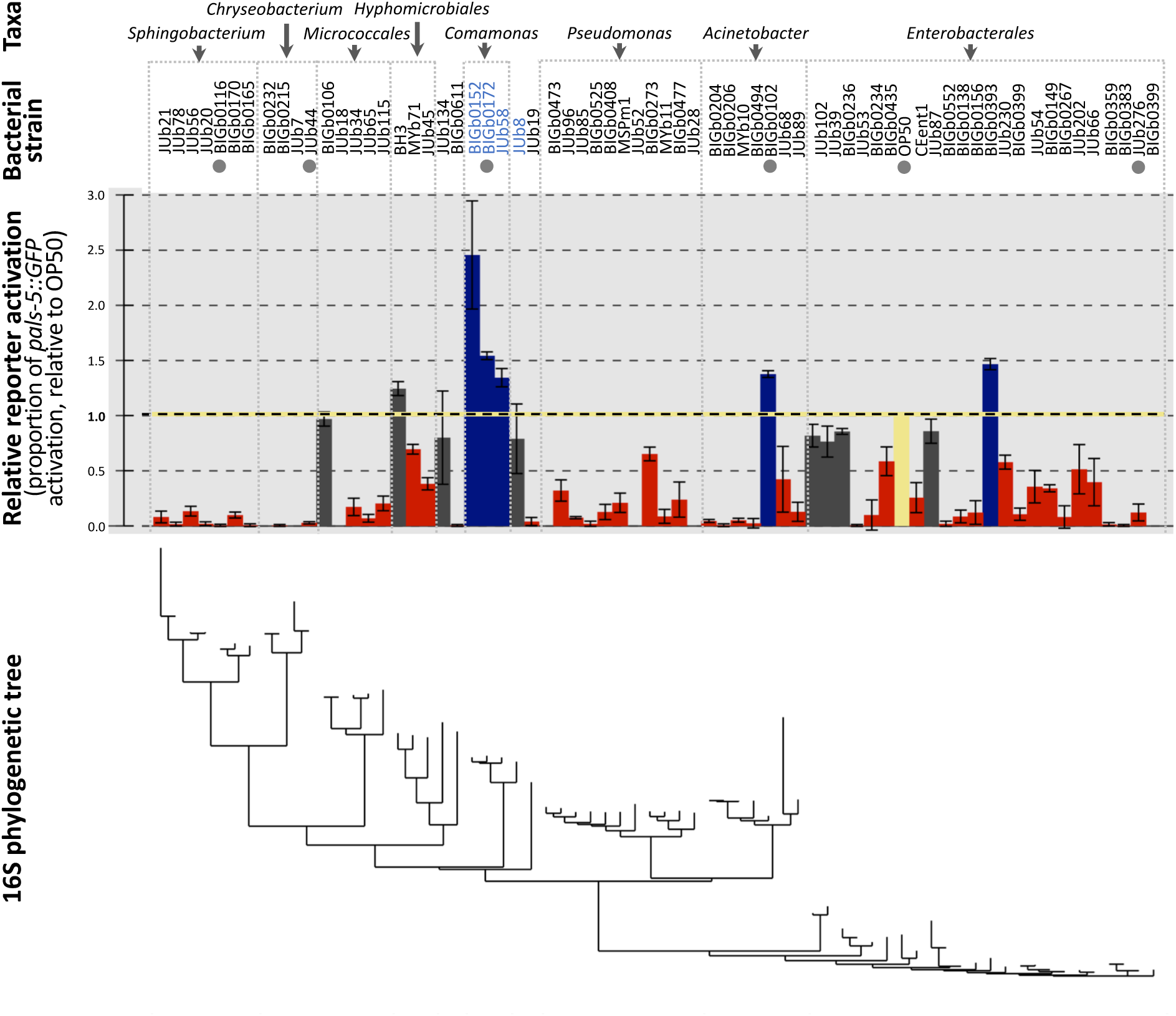
Most natural bacteria reduce *C. elegans* susceptibility to viral infection. For each bacterial environment, three independent populations of ≍100 animals were challenged with virus and the proportion of animals activating the *pals-5* reporter was measured. Data were collected in three experiments and the values for each bacteria are shown relative to the values of *Escherichia* OP50 in the corresponding experiment. The *Escherichia* OP50 reference bar is highlighted in yellow, while bacterial strains that induce a significant reduction of infection are colored in red, those with no significant differences with OP50 in grey, and strains with a significantly higher activation of the *pals-5* reporter are displayed in blue. Data are presented as mean ± standard error. Bacteria are arranged according to their phylogenetic relationships, with taxonomic classifications provided in the top rows and a phylogenetic tree based on their 16S sequences in the bottom row. Strain names in blue indicate a significant phylogenetic signal for these strains. Natural strains marked with a grey dot were selected for further investigation.

Our screen includes a phylogenetic diverse set of 67 bacterial strains (Figure 1). Using the Local Indicator of Phylogenetic Association (local Moran’s I test), we detected no phylogenetic signal for most strains, suggesting that the trait values are randomly distributed across the tree at this level of sampling. However, we identified a significant phylogenetic signal in *Comamonas* strains plus *Delftia* (*P* < 0.01), indicating a potential phylogenetic clustering for these four strains in triggering a particularly high *C. elegans* response to viral infection. On the other side of the spectrum, we note that all eleven tested members of Bacteroidetes completely suppressed *pals-5* expression upon OrV exposure.

In four cases, our screen included two distinct strains labeled with the same species name, indicating a particularly close relationship. The different strains of each species have similar effects expect for those of *Leucobacter luti* BIG0106 and JUb18 (Supplementary Figure 3A, B). We also tested various *E. coli* strains and found that despite some small but significant differences, all enabled a similar level of activation of the *pals-5* reporter in virus-inoculated animals (Supplementary Figure 3C).

This initial screen was performed using *pals-5* as an infection reporter. To assess more directly the effect of the bacterial environment on viral infection, we directly stained viral RNA using fluorescence in situ hybridization (FISH). We focused on five bacteria (indicated by grey dots in Figure 1): two enhancers and three suppressors in the *pals-5* screen. On the enhancing *Acinetobacter* BIGb0102 and on the three suppressive bacteria, the viral FISH staining matched the result with the *pals-5* reporter (Figure 2A-B). However, in the *Comamonas* BIGb0172 environment, the proportion of FISH-stained animals remained similar to that in OP50 (Figure 2A). We further tested on the two enhancing bacteria fluorescent reporters for the *F26F2.1* or *sdz-6* genes that are part of the IPR response but, unlike *pals-5*, are not controlled by the transcription factor ZIP-1 (33). For *Acinetobacter* BIGb0102 we observed a similar pattern of gene activation for *pals-5* than for *F26F2.1* and *sdz-6*. However, in the *Comamonas* BIGb0172 environment, only *F26F2.1* activation was higher than in OP50 upon viral infection (Figure 2C). In conclusion, the *Acinetobacter* BIGb0102 environment enhances viral infection compared to *Escherichia* OP50 while *Chryseobacterium* JUb44, *Sphingobacterium* BIG0116, and *Lelliottia* JUb276 strongly suppressed the infection level.

**Figure 2.**
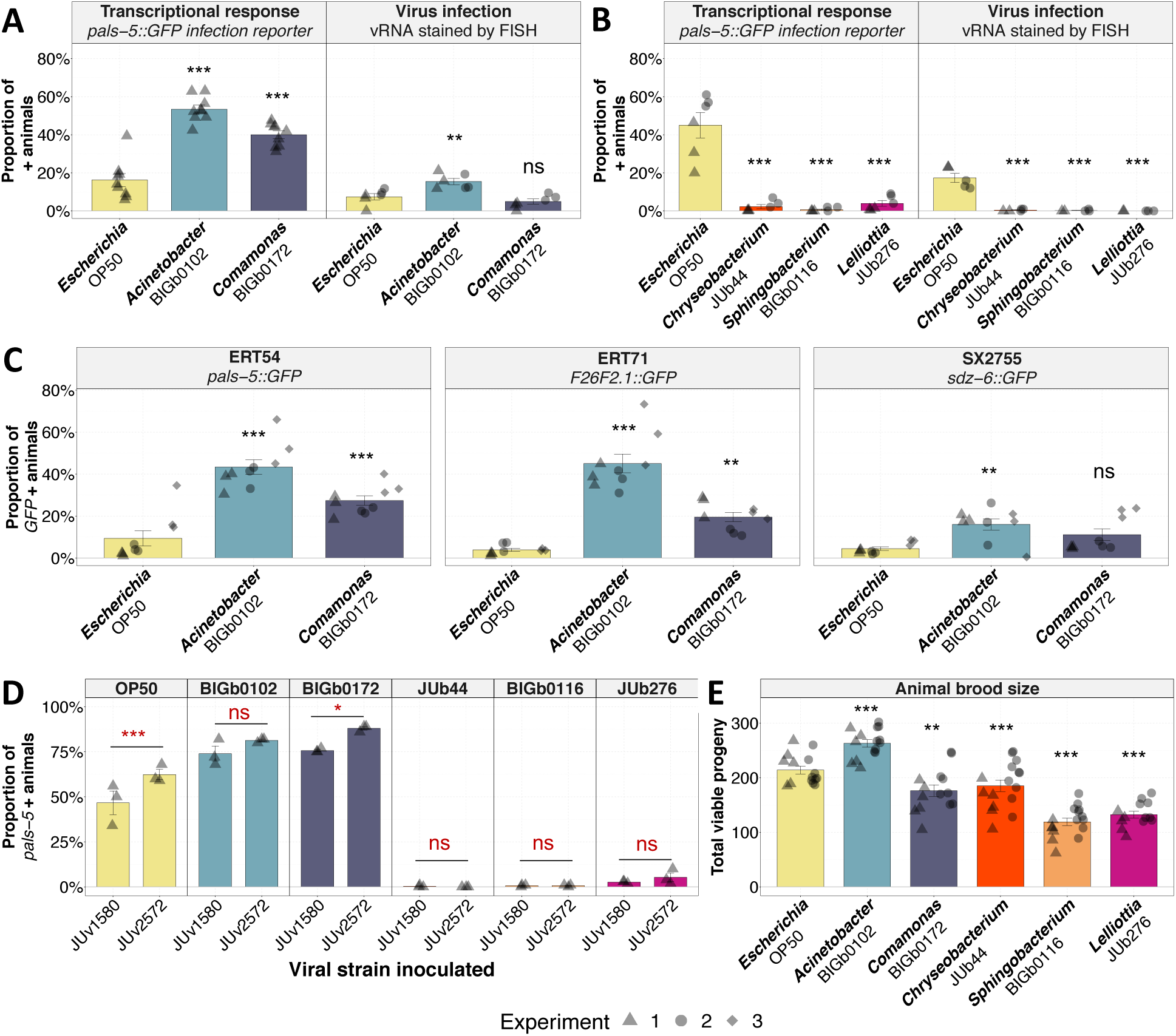
Study of selected natural strains confirms the bacterial modulation of nematodes susceptibility to viral infections **(A-B)** Transcriptional response and proportion of infected animals for bacteria that induced (A) strong activation or (B) no activation of pals-5 in the initial screen. **(C)** Activation of three intracellularpathogen transcriptional response genes (*pals-5*, *F26F2.1*, and *sdz-6*) in animals exposed to virus in *Acinetobacter* BIGb0102 or *Comamonas* BIGb0172. **(D)** Transcriptional response of animals exposed to natural bacteria when challenged with a more infective Orsay virus strain. **(E)** Total viable progeny of ERT54 animals cultured on the selected bacteria. Each data point represents an independent population of ≍100 animals. Black symbols indicate the significance of the difference between the labeled bacteria and the *Escherichia* OP50 reference; Red symbols indicate the significance of the difference between treatments on the same bacteria. Data are presented as mean ± standard error. Asterisks on the graphs represent values of significance: *** P < 0.001; ** P < 0.01; * 0.01 < P < 0.05; P values higher than 0.05 are labeled as “ns” (same symbols for all the figures of this study).

**Figure 3.**
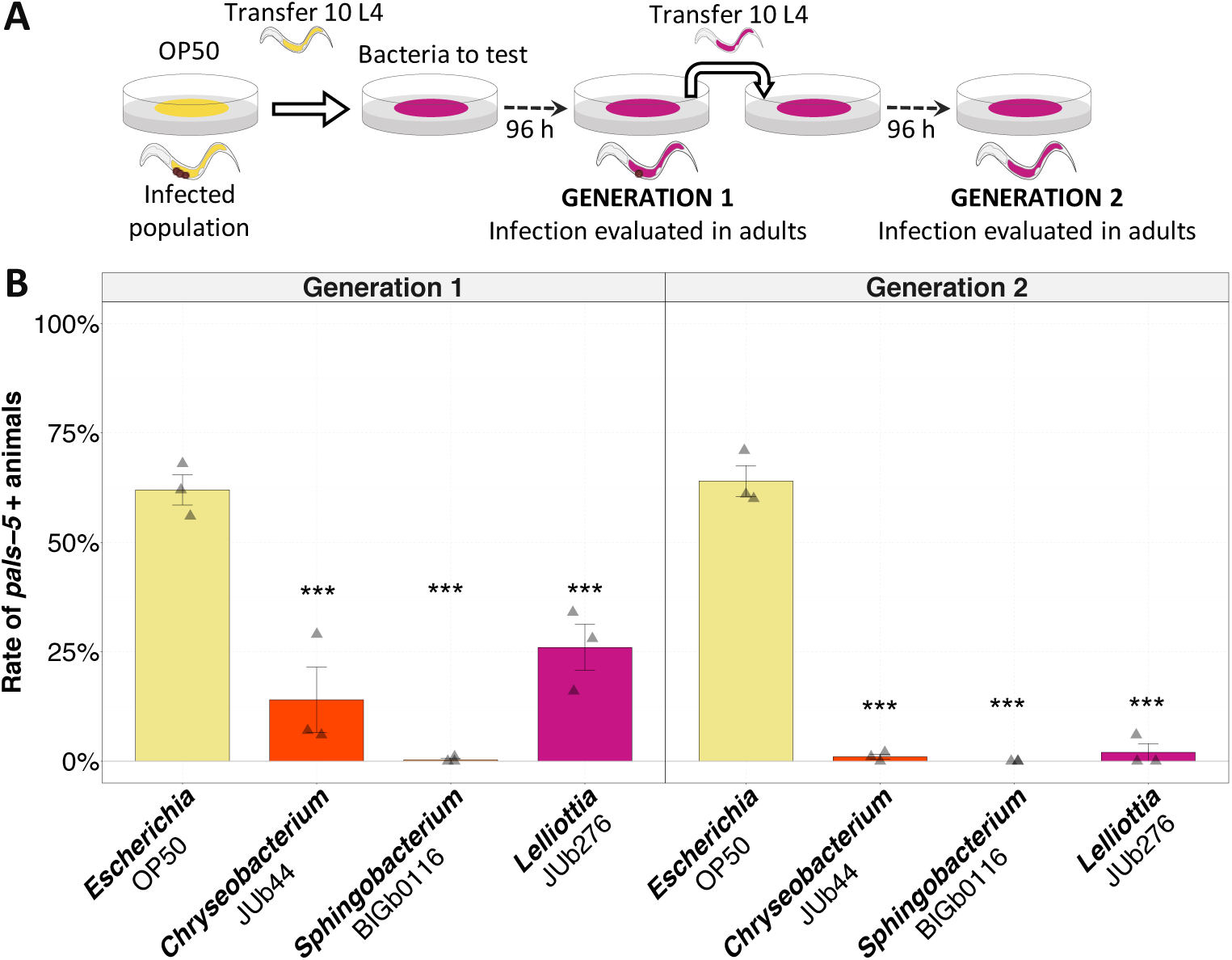
Natural bacteria suppress the OrV persistence over generations. **(A)** Schematic representation of the experimental design. **(B)** Activation of the *pals-5* reporter in animals transferred to different natural bacteria at two successive generations.

We tested whether the effect of bacterial environments was specific to the conventional viral strain used in our main experiments (ORV strain JUv1580). We tested JUv2572, an OrV strain reported by Frézal et al. (39) to be more infectious than JUv1580 and prone to infecting more anterior host intestinal cells. Resistance-inducing bacteria induced resistance even with this stronger viral strain (Figure 2D).

We wondered whether the bacteria favoring viral infection did so by generally weakening the host compared to those suppressing it. We thus measured the production of offspring over time on each of the five bacteria and *E. coli* (in the absence of virus) and found that it could not be the case. Contrary to the expectation if BIGb0102 weakened *C. elegans*, animals in the enhancing *Acinetobacter* BIGb0102 had a significant increase in total brood size compared to *Escherichia* OP50 (Figure 2E), while instead *Comamonas* BIGb017, *Lelliottia* JUb276, and *Sphingobacterium* BIGb0116 caused a significant decrease. Except for *Acinetobacter* BIGb0102, all bacterial environments significantly delayed the production of offspring compared to *Escherichia* OP50 (Supplementary Figure 4), that is, there was no significant difference between the resistant-inducing *Chryseobacterium* JUb44 and the susceptibility-inducing *Comamonas* BIGb0172. We thus concluded that the bacterial environments that enable a strong viral infection did not do so by generally weakening the host.

### Natural bacteria eliminate OrV in infected nematode populations within two generations

In natural environments, viral infections spread within populations as infected organisms continuously produce and release viruses, which facilitates transmission to other susceptible hosts (40). This differs from our experimental setup, which involves a single viral input, limiting the potential for continuous spread and transmission of the virus within the tested population. In order to create conditions of horizontal infection from other individuals, we transferred individuals from a previously infected *C. elegans* population on *Escherichia* OP50 to resistance-inducing bacterial environments (Figure 3). We found that the offspring of nematodes transferred to a resistance-inducing bacterial lawn had reduced activation of the *pals-5* infection reporter compared to those transferred to *Escherichia* OP50. Activation of *pals-5* was almost absent in the second generation of nematodes cultured in all the tested bacteria, except for the control (*Escherichia* OP50), which showed similar levels to the first generation (Figure 7B). We observed the same pattern of virus elimination in the second generation when the transfer was performed by transferring a piece of agar with a random number of nematodes from an old plate to a new plate (Supplementary Figure 5A).

### Resistance-inducing bacteria do not reduce the virion infectivity

To examine whether the bacterial lawn affected virion infectivity prior to entering the host, we employed two approaches: (i) pre-incubating the virus with bacteria before inoculation, and (ii) inoculating the nematodes with the virus in *Escherichia* OP50 and then transferring them to another bacterial environment.

In the first approach, we incubated the virus with either *E. coli* or the natural bacteria at 20°C for 24 h (Figure 4A). After incubation, bacteria were pelleted and the filtered viral supernatants were inoculated onto animals in *Escherichia* OP50 environments. We did not see a significant difference in the final infectivity of the viral preparations (Figure 4A).

**Figure 4.**
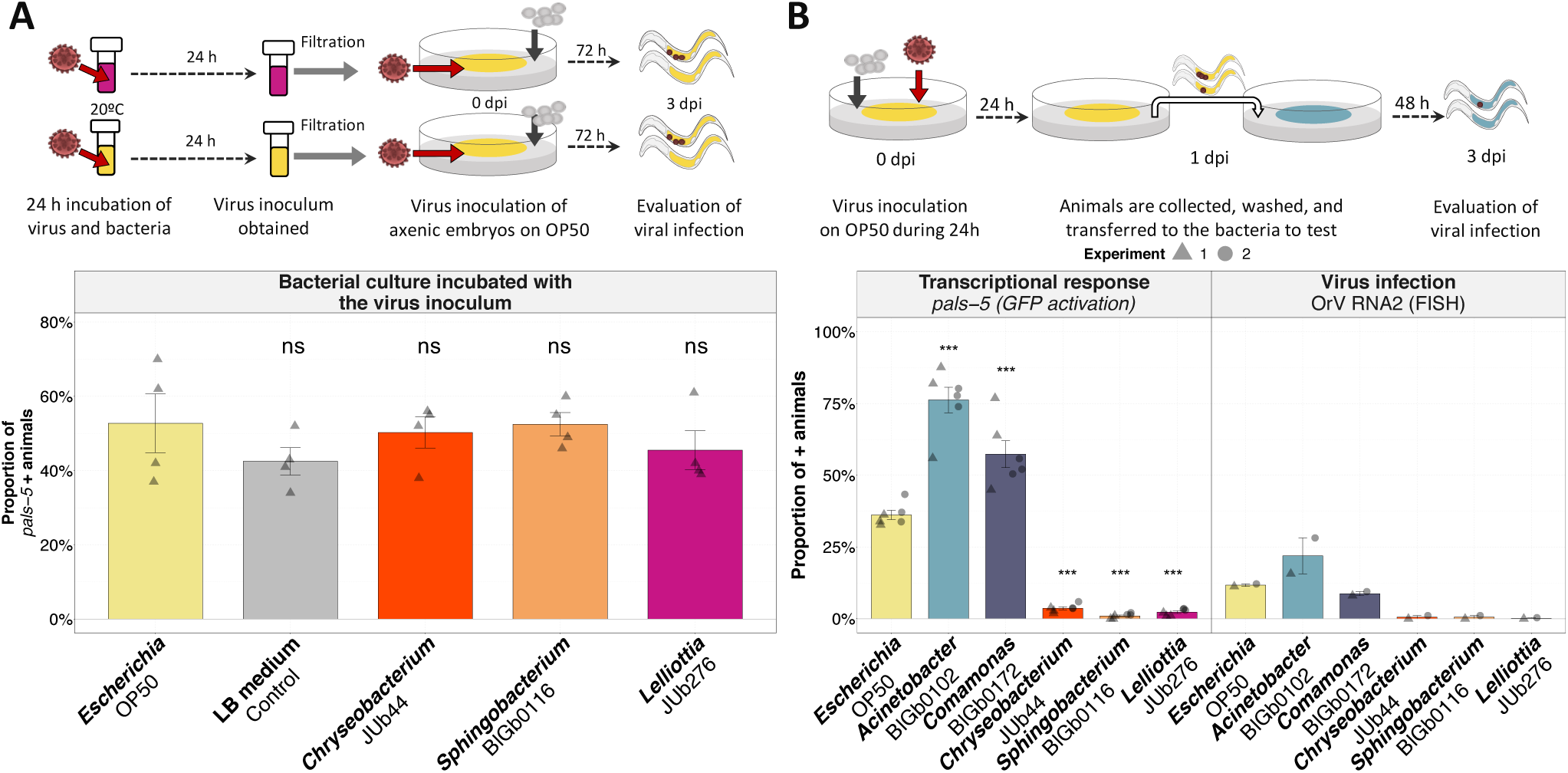
Assessment of potential virion degradation by different bacteria. **(A)** Activation of an infection reporter in animals on *Escherichia* OP50, challenged with viruses previously incubated with various bacterial cultures. **(B)** Animal response to OrV challenge after initial exposure to the virus in *Escherichia* OP50 and subsequent transfer to different, non-virus-inoculated, bacteria.

**Figure 5.**
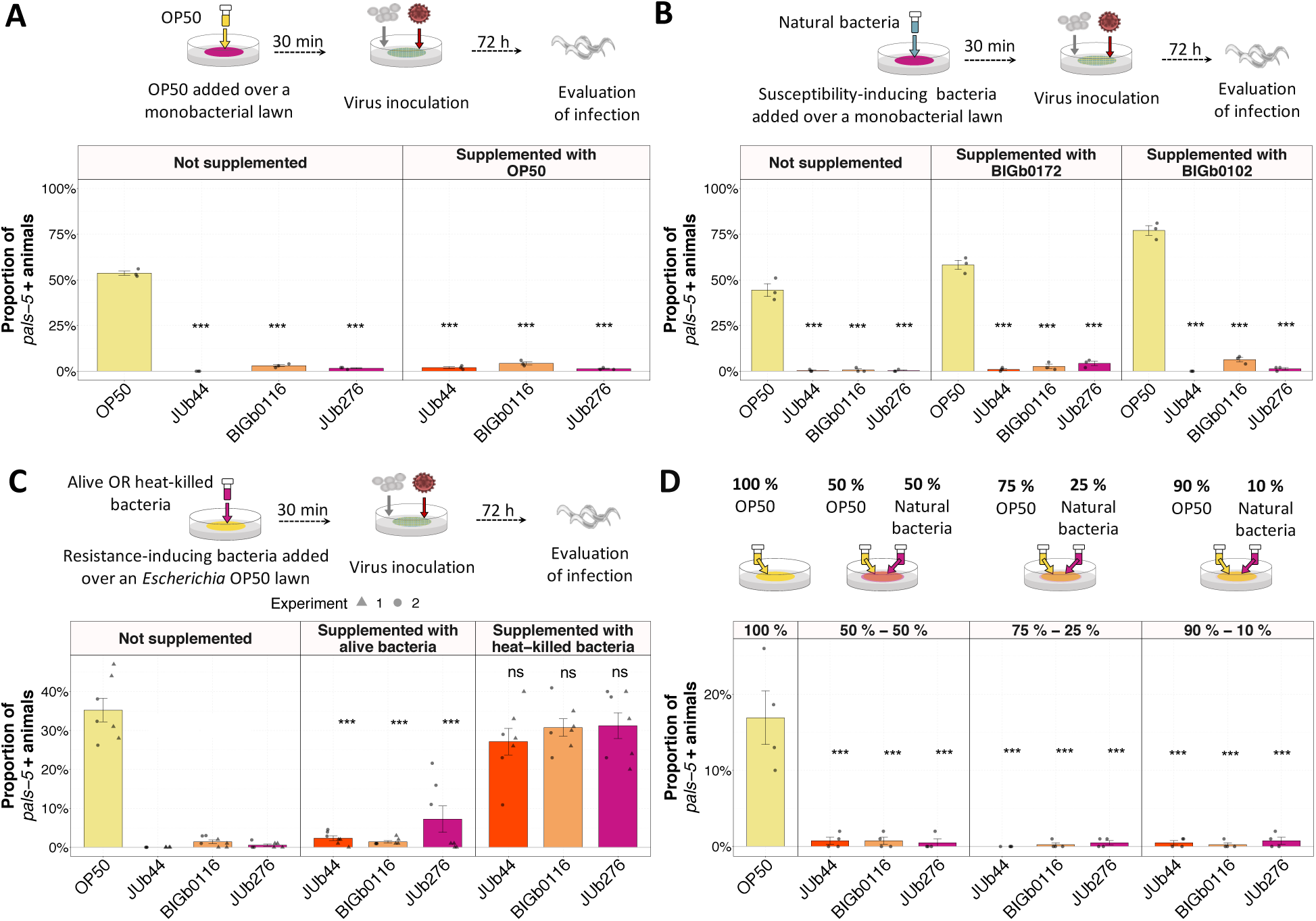
Resistance-inducing bacteria can induce resistance in mixed bacterial environments, but this effect is not seen if the bacteria are heat-killed. **(A-D)** Activation of *pals-5* reporter in animals on monobacterial environments (not-supplemented) or (A) on natural bacteria supplemented with *Escherichia* OP50, (B) on natural bacteria supplemented with natural bacteria reported to enhance viral infection, (C) on *Escherichia* OP50 supplemented with live or heat-killed cultures of resistance-inducing natural bacteria, or (D) on bacterial lawns seeded with mixed combinations of bacteria. In all panels, the top row shows a schematic representation of the experimental design.

In the second approach, we exposed axenic embryos to OrV for 24 hours on *Escherichia* OP50 lawns and then transferred the larvae to plates with natural bacterial lawns (Figure 4B). These results indicate that the natural bacteria do not suppress by degrading the virions outside the host; natural bacteria can alter transcriptional response and infection susceptibility after initial virus exposure in a susceptible environment.

### Resistance-inducing bacteria effect prevails in mixed bacterial environments

We then wondered whether a suboptimal diet could be the cause of viral resistance. We supplemented natural bacterial lawns with bacteria that enable viral infection. To minimize potential interactions and competition between the bacteria, we prepared plates with the natural bacteria and added permissive bacteria: *Escherichia* OP50 (Figure 5A) or *Acinetobacter* BIGb0102 and *Comamonas* BIGb0172 (Figure 5B), right before transferring axenic embryos and inoculating OrV. Despite the supplementation, *Chryseobacterium* JUb44, *Sphingobacterium* BIGb0116, and *Lelliottia* JUb276 induced resistance to infection. When we added resistance-inducing natural bacteria onto *Escherichia* OP50, resistance induction persisted. However, this effect was abolished when the natural bacteria were first heat-killed (Figure 5C). These three natural bacteria induced resistance to viral infection even in mixed environments (with *Escherichia* OP5O) where their initial presence was as low as 10% (Figure 5D).

The prevalence of resistance-inducing effects was also observed when testing the CeMbio community, a defined natural and ecologically relevant bacterial community (41). Individual strains in the CeMbio community had diverse effects on susceptibility to infection. However, nematodes were resistant to infection when associated with the CeMbio community (Supplementary Figure 6D). Thus, bacterial-induced resistance prevails over permissive bacteria. These findings thus reject the poor diet hypothesis.

**Figure 6.**
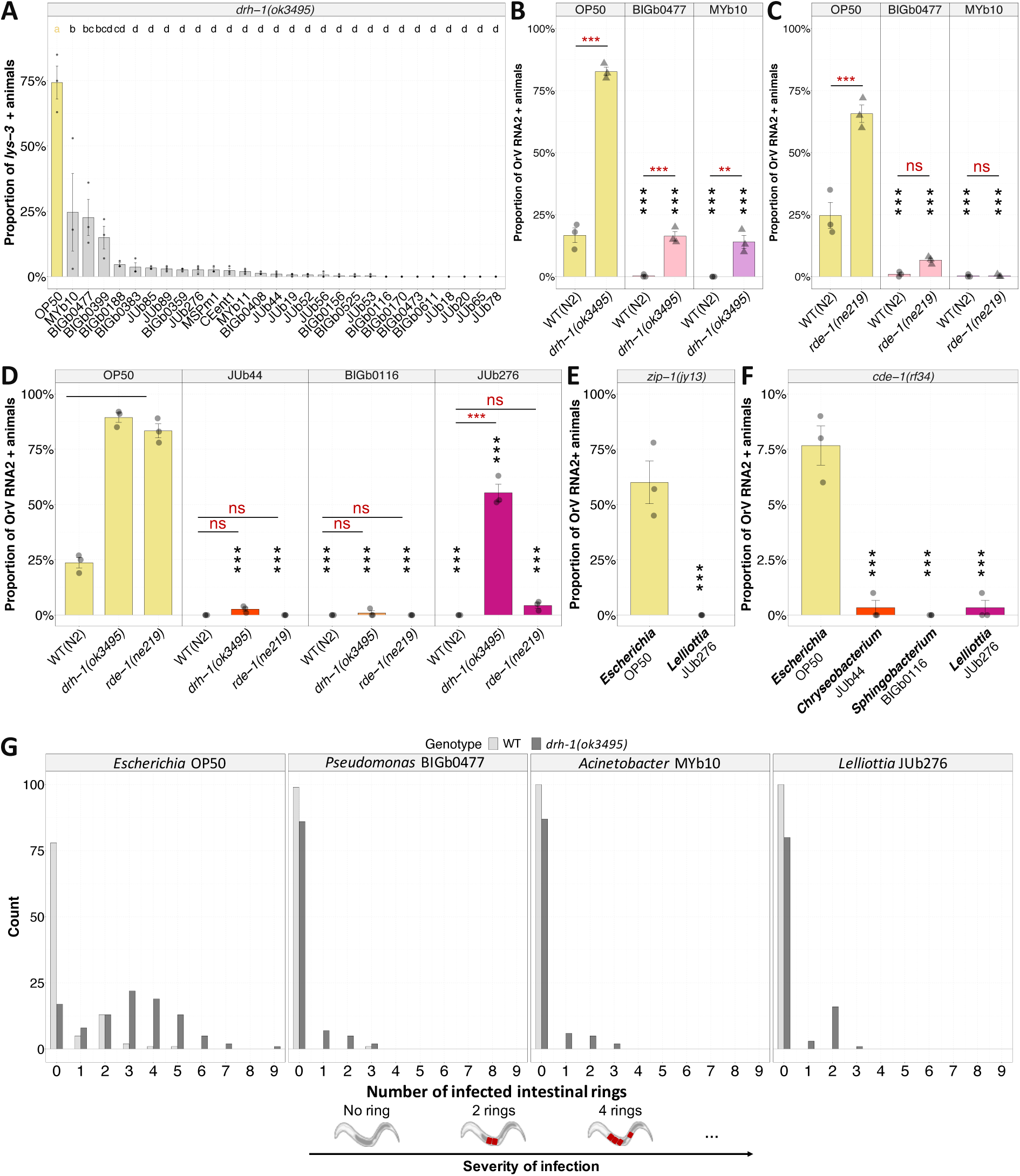
Susceptibility to OrV infection in animals with hampered antiviral immune responses on resistanceinducing bacterial environments. **(A)** Proportion of *drh-1* animals on different OrV resistance-inducing bacteria that activate the *lys-3* reporter after being OrV challenged. **(B-C)** Proportion of infected (B) WT and *drh-1* or (C) WT and *rde-1* animals in bacteria that enable slight activation of the *lys-3* reporter in panel A. **(D-G)** Proportion of infected animals with hampered antiviral immune responses on selected bacteria: (D) *drh-1* and *rde-1*, (E) *zip-1*, and (F) *cde-1*. **(G)** Spread of viral infection within the intestinal ring of the nematode, which serves as a proxy for infection severity, for WT and *drh-1* animals.

### Unknown antiviral pathways are involved in the bacterial suppression of viral infection

We sought to identify host pathways involved in the suppressive effect of bacteria on viral infection. To accomplish this, we evaluated viral infection in animals carrying null alleles for key components of the antiviral response. Antiviral immunity starts with DRH-1/RIG-I triggering both the small RNA response and the IPR transcriptional response (therefore lowering the ubiquitin-mediated immunity). We thus tested whether the bacteria suppressed infection in the *drh-1* mutant context. As the IPR response is not activated in *drh-1* mutants, we could not use *pals-5* as the reporter and instead utilized a *lys-3* reporter, which is activated upon severe infection (34). 75 % of *drh-1* animals activated the *lys-3* reporter in the *Escherichia* OP50 environment, however the *lys-3* reporter was barely activated in the suppressive natural bacteria compared to *Escherichia* OP50 (Figure 6A). Thus, the suppression is mostly independent of DRH-1. In three bacteria (*Raoultella* BIGb0399, *Pseudomonas* BIGb0477, and *Acinetobacter* MYb10), the activation of *lys-3* was less strongly reduced (Figure 6A). We chose two of the latter bacteria (Figure 6B,C) and the three suppressive bacteria studied above (Figure 6D-E) and repeated the experiment, comparing the level of viral infection in wild-type and *drh-1* mutant animals using viral RNA FISH. In the *Chryseobacterium* JUb44 and *Sphingobacterium* BIGb0116 environments, the virus could not infect even the *drh-1* mutant (Figure 6D), indicating that these bacteria do not require the DRH-1 dependent pathways to repress OrV infection.

In the *Pseudomonas* BIGb0477, *Acinetobacter* MYb10, and *Lelliottia* JUb276 environments, the *drh-1* animals showed a lower infection than on *Escherichia* OP50, but significantly increased infection compared to the wild-type animals (Figure 6B,D; Supplementary Figure 5B). We confirmed the results using an alternative *drh-1* allele mimicking a natural deletion (31) (Supplementary Figure 6C). The severity of the infection was low in these FISH-stained animals (Figure 6H). Thus, even for these bacteria, a part of the suppressive effect is also DRH-1 independent.

Downstream of DRH-1, we tested mutants in RDE-1, the Argonaute required for the RNAi interference response, and in the ZIP-1 transcription factor required for part of the DRH-1 dependent transcriptional response (Figure 6C,E,F). In these mutants, the suppressive bacteria, including *Lelliottia* JUb276, fully suppressed infection. Thus, the weak infection observed in *drh-1* mutants on *Lelliottia* JUb276 does not seem to be RDE-1 or ZIP-1 dependent.

We tested *cde-1* mutants, which are defective in the viral uridylation response. *cde-1* mutants are resistant to viral infection in *Chryseobacterium* JUb44, *Sphingobacterium* BIGb0166, and *Lelliottia* JUb276 environments (Figure 7G). To test the possibility that immune pathways against bacteria, such as p38/PMK-1 may be involved in the induced resistance against viruses, we tested some mutants on the antibacterial immune response. The *Lelliottia* JUb276 environment also induced resistance to the virus in these mutants (Supplementary Figure 7).

In conclusion, for most suppressive bacteria, the reduction in viral infection is independent of known antiviral responses including RNA interference, transcriptional regulation downstream of DRH-1 and uridylation. However, *Pseudomonas* BIGb0477, *Acinetobacter* MYb10, and *Lelliottia* JUb276 fail to induce resistance to viral infection when DRH-1 is not functional.

## DISCUSSION

We conducted a comprehensive study examining the impact of naturally associated bacteria on *C. elegans* susceptibility to viral infections. Our screen involved monocultures of 67 bacteria, revealing that monobacterial environments significantly influence host susceptibility to viral infections, with some bacteria providing protective effects, while others increase the host’s susceptibility. Notably, the majority of natural bacteria tested reduced host susceptibility to viral infection compared to the non-natural bacterial environment commonly used in research. This could explain the difficulties for isolating natural viruses (27) and highlights the importance of considering the laboratory microbial environment when isolating natural strains for the study of their viruses. In our screen, strains of the same species generally exhibited similar effects on host resistance but two geographically distant *Leucobacter luti* strains (JUb18 from South Africa and BIGb0106 from France; 15) exhibited distinct effects on host resistance to infection. Investigating differences between such strains or conducting a genetic screen in one of them could provide valuable insights into the bacterial factors involved in resistance to viral infections.

Defensive microbes can provoke fitness consequences for their hosts (42). The trade-offs of the induced resistance are essential in understanding the complex interplay of host-microbes interactions. We found that certain bacterial environments may confer protection against viral infections while simultaneously reducing host fitness. In contrast, other environments may achieve a more optimal balance between defense against viral infections and maintenance of host fitness. Gaining insights into these trade-offs is pivotal for devising effective strategies to bolster host immunity while mitigating potential adverse effects on other phenotypes of interest.

Host nutrition can have diverse effects in pathogens (43). In *C. elegans*’ bacterial environments there is no distinction between food (nutrition) and biotic environment. This overlap is likely significant, given that the lipid content of the nematodes plays a crucial role in viral infections (44). Our experiments suggest that the resistance to virus infection is not caused by nutritional deficits because supplementation with bacteria that enable virus infection does not recover susceptibility to the virus. Some bacteria have been shown to have immunomodulatory properties even after being heat-killed (45). Our tested bacteria do not induce resistance after being heat-killed, indicating that either the bacteria need to be alive to induce resistance or that the bacterial factor that induces resistance is thermolabile.

We found that some bacteria can induce resistance to infection without requiring known host antiviral mechanisms. Some bacteria fail to induce resistance in *drh-1* mutants but could induce it in mutants of known DRH-1 downstream pathways. DRH-1 recognizes the viral genome as foreign, possibly through its well-conserved RIG-I domain (31), and activates multiple immune responses (reviewed in 29) and transcription of genes of unknown function. Our findings suggest that either some bacteria fail to induce resistance in the highly susceptible *drh-1* mutants or that the bacteria instigate resistance partially through DRH-1, via an unknown downstream pathway.

Our research emphasizes the importance of considering the natural biotic environment in the study of viral infections and their ecological and evolutionary implications (46, 47). This study provides a valuable foundation for future research by suggesting the involvement of unknown antiviral mechanisms and by establishing a basis for dissecting the molecular mechanisms underlying host-virus interactions accounting for the biotic environment. Future research should aim to uncover the molecular mechanisms underlying these protective effects and explore their applicability to a broader range of viral strains and host species. This understanding may help to discover unknown immune mechanisms, develop new strategies for modulating host susceptibility to viral infections and improving host health through targeted manipulation of the microbes associated with an organism.

## MATERIAL AND METHODS

### *C. elegans* strains and maintenance

Nematodes were cultured on Nematode Growth Media (NGM) plates at 20°C, following standard procedures (48, 49). The NGM plates were seeded with 100 µL of a monobacterial culture. The nematode strains used in this study are listed in Supplementary Table 2. The *drh-1(mcp553)* allele mimics the natural deletion in JU1580 and other *C. elegans* wild isolates (31). It was created using a CRISPR/Cas9-mediated edition in the N2 background by the CNRS Segicel Platform (Lyon, France) using two guides crMG023 (gCTATCGTGTTGCTAGTCGA)and crMG024 (ACCGACCGAAATACGACAAT) and the repair template (tctttacatgcttattttatttaattcttaattctattaattatttaattttcagctatc AATGAGAGATGCGGATCAAGCTCGAACACCAATGGTATTTGAGCATCACGCGAATGG AGA).

### Bacterial maintenance

Bacteria were cryo-preserved at -80°C in Luria broth medium (LB; 10 g/L tryptone, 5 g/L yeast extract, 5 g/L NaCl) with 25 % glycerol. For cultivation, a cryo-preserved culture was streaked on LB-agar plates and incubated at room temperature for 48 h. Colonies were then picked and inoculated into 5 mL of liquid LB. The culture was grown at 28°C and 220 rpm for 16 h for the natural strains, while *E. coli* OP50 was grown at 37°C. The bacterial strains used in this study are listed in Supplementary Table 1. Bacteria were heat-killed by placing the bacterial cultures in a water-bath at 100°C for 40 min.

### Virus maintenance

The Orsay virus strain JUv1580 (Félix et al., 2011) was used in all experiments except when indicated, where the strain JUv2572 was used (39). Viral preparations were prepared by inoculating a viral inoculum derived from the original JU1580 infected animals (26) on 90 mm OP50-seeded plates with JU2624 animals (*C. elegans* JU1580 isolate in which the *lys-3::eGFP* construct was introgressed by 10 rounds of backcrosses to JU1580; 34). The animals were collected with M9 buffer four days after inoculation, pelleted, and the supernatant was filtered through a 0.22 µm filter to obtain the OrV-infectious supernatant. To create a stock of the viral inoculum, the supernatant was aliquoted and cryo-preserved at -80°C.

### Virus inoculation procedure

To synchronize the nematode population and obtain axenic animals, we treated young adult populations with 4 mL of a sodium hydroxide and sodium hypochlorite mixture (2 mL sodium hypochlorite 12%, 1.25 mL NaOH 10 N, 6.75 mL H2O) for three minutes. Subsequently, we washed the samples four times with 15 mL of M9 buffer. This procedure resulted in axenic embryos, which were placed around the bacterial lawn previously inoculated with 50 µL of the viral inoculum. The inoculated plates with animals were maintained at 20°C, and infection was assessed 72 hours post-inoculation.

### Evaluation of viral infection

The activation of the intracellular pathogen response was measured using a fluorescent reporter activated upon intracellular infection (*pals-5::GFP*; 38) or a reporter of severe biotic stress (*lys- 3::eGFP*; Le Pen et al., 2018). The proportion of infected animals was analyzed by visualizing viral RNA on nematodes intestinal cells; the OrV RNA was stained using fluorescent in situ hybridization (FISH), as described by Frézal and colleagues (39). In brief, animals were collected and fixed in a 10% formamide solution. Fixed worms were stained targeting OrV RNA2 molecules using a 1:40 dilution of a mix of oligonucleotide sequences (39, 50) conjugated to the Cal Fluor red 610 fluorophore or a non-diluted single probe (5’ ACC ATG CGA GCA TTC TGA ACG TCA 3’) conjugated to Texas Red. Stained animals were examined using a Zeiss AxioImager M1 compound microscope with 10× (0.3 numerical aperture) and 40× (1.3 numerical aperture) objectives. An animal was considered infected if at least one intestinal cell displayed distinct fluorescence at higher levels than the background staining for that individual.

### Phylogenetic analysis

The 16S sequences used in the phylogenetic analysis can be found in Supplementary File 1. A multiple sequence alignment of the sequences was performed using the MAFFT v7 tool (51) of MPI Bioinformatics Toolkit (52). Aligned sequences were used to infer the phylogenetic tree by maximum likelihood using the IQ-TREE web server (53). The phylogenetic tree and the plot of the Figure 1 were generated using the *phylo4d* function from the “phylosignal” package version 1.3 (54). The phylogenetic signal in the trait data was assessed using the *lipaMoran* function, also available in the “phylosignal” package. To account for multiple testing, we adjusted the obtained p-values using the Benjamini-Hochberg method. The level of significance was set at *P* < 0.01.

### Statistical analysis

All statistical analyses were conducted using R version 3.6.1 within the RStudio development environment version 1.3.1093. The level of significance was set at *P* < 0.05. The significance of the differences between groups were calculated using:

Figure 1: an analysis of variance and Dunnett’s contrasts using *Escherichia* OP50 as the reference group.

Figure 2A,B,C,E: a linear-mixed model, where the factor was the bacteria and the experiment were considered a random effect. Post hoc multiple comparisons were performed using Tukey contrasts.

Figure 2D: a generalized linear model with logistic regression, where the factors were bacterial and viral strain. Post hoc multiple comparisons were performed using Tukey contrasts.

Figure 3: a generalized linear model with logistic regression, where the factors were bacteria and generation. Post hoc multiple comparisons were performed using Tukey contrasts.

Figure 4A: using analysis of variance with bacteria as a factor. Post hoc multiple comparisons were performed using Tukey contrasts.

Figure 4B: a linear-mixed model, where the factor was the bacterial environment and the experiment was considered a random effect.

Figure 5A,B,C: a generalized linear model with logistic regression, where the factors were bacteria and treatment. Post hoc multiple comparisons were performed using Tukey contrasts.

Figure 5D: a linear-mixed model, where the factors were the bacteria and the treatment, and the experiment was considered a random effect. Post hoc multiple comparisons were performed using Tukey contrasts.

Figure 6A,E,F: an analysis of variance with bacteria as a factor. Post hoc multiple comparisons were performed using Tukey contrasts.

Figure 6B,C,D: a generalized linear model with logistic regression, where the factors were bacteria and host genotype. Post hoc multiple comparisons were performed using Tukey contrasts.

The analysis of variance, Tukey or Dunnett’s post hoc test, and letter-based grouping for multiple comparisons were performed using the following functions: (i) the *aov()* function from the “stats” package version 3.6.1 (55) was used for conducting the ANOVA (ii) the *TukeyHSD()* function from the ’stats’ package was used for performing the Tukey post hoc test (iii) Dunnett’s post hoc test was carried out using the *glht()* function from the “multcomp” package version 1.4-23 (56) (iv) the *multcompLetters4* function from the “multcompView” package version 0.1.8 (57) was used for generating letter-based grouping for multiple comparisons. Generalized linear models were performed using the *glm* function from the “stats” package. To account for the block effect, we employed a linear mixed-effects model. The model was fitted to the data using the *lmer* function from the “lme4” package version 1.1-21 (58). Post hoc pairwise comparisons were done using the *emmeans* function from the “emmeans” package version 1.4.7 (59). Tukey’s method was used to adjust for multiple comparisons in the pairwise comparisons.

Graphs were generated using the “ggpubr” package version 0.2.4 (60) in R. Data are presented as mean ± standard error. Asterisks on the graphs represent values of significance: *** *P* < 0.001; ** *P* < 0.01; * 0.01 < *P* < 0.05; *P* values higher than 0.05 are labeled as *ns*.

## DATA AVAILABILITY

All data generated in this study can be found in Supplementary File 2.

## ACKNOWLEDGEMENTS

We thank Aurélien Richaud for excellent technical assistance and advice and Tony Bélicard for the JU2624 strain. We want to thank Emily Troemel for sharing the *zip-1(jy13)* strain. Some strains were provided by the CGC, which is funded by NIH Office of Research Infrastructure Programs (P40 OD010440). We thank the CNRS Segicel platform for generating the *drh-1(mcp553)* strain. We thank WormBase. We also thank members of Marie-Anne Félix, Marie Gendrel, and Henrique Teotónio teams for fruitful discussions. R.G. is funded by an EMBO Postdoctoral Fellowship (ALTF 311-2021). M.A.F. is funded by the Centre National de la Recherche Scientifique. This work was partially funded through a grant from the Agence Nationale de la Recherche ANR-19-CE12- 0025.

## SUPPLEMENTARY MATERIAL

**Supplementary Figure 1.**
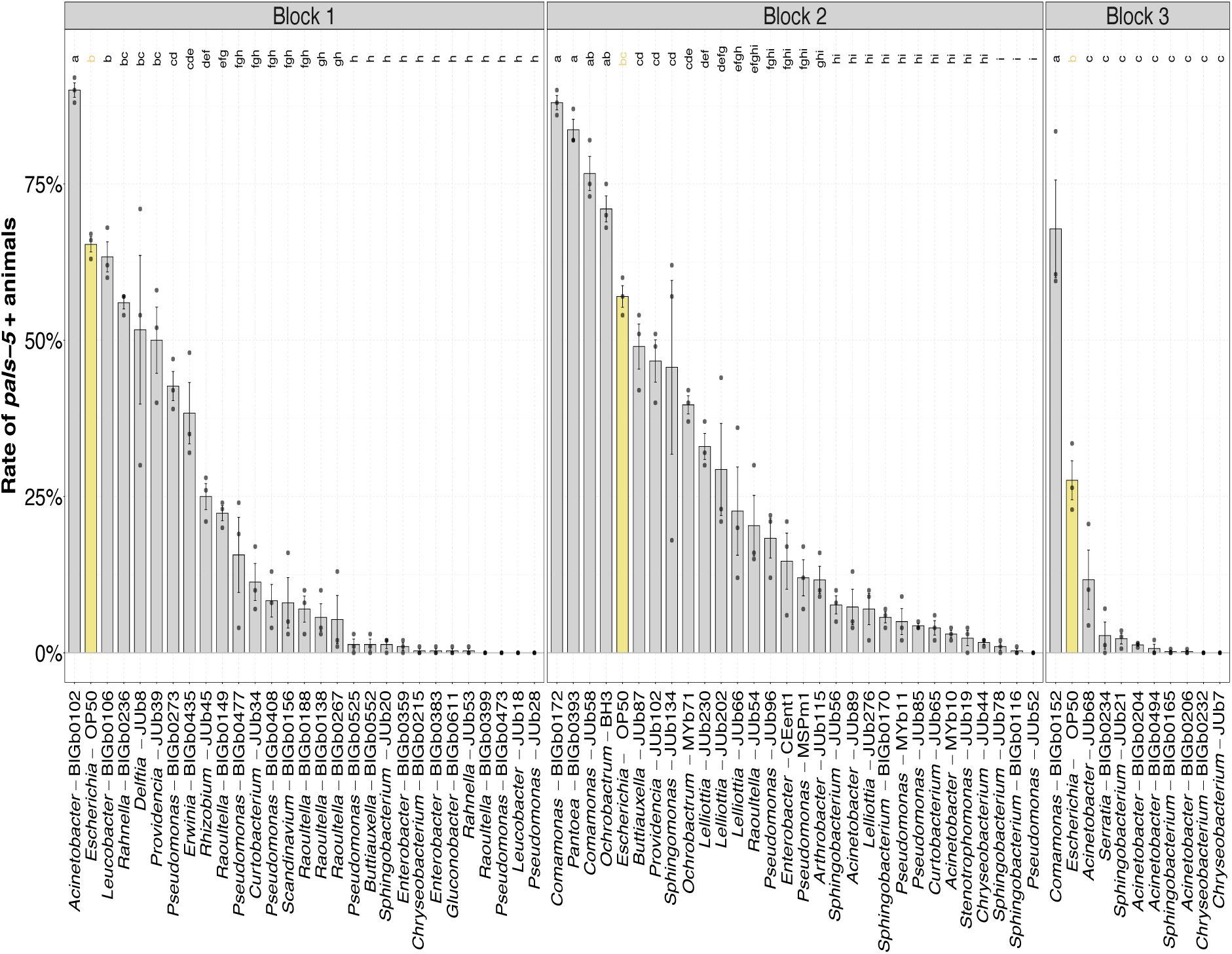
Proportion of animals with activation of the *pals-5* reporter after being challenged with virus; original data for each block, with 3 replicates per bacteria. Letters over the bars indicate letter-based grouping for multiple comparisons. Data are presented as mean ± standard error.

**Supplementary Figure 2.**
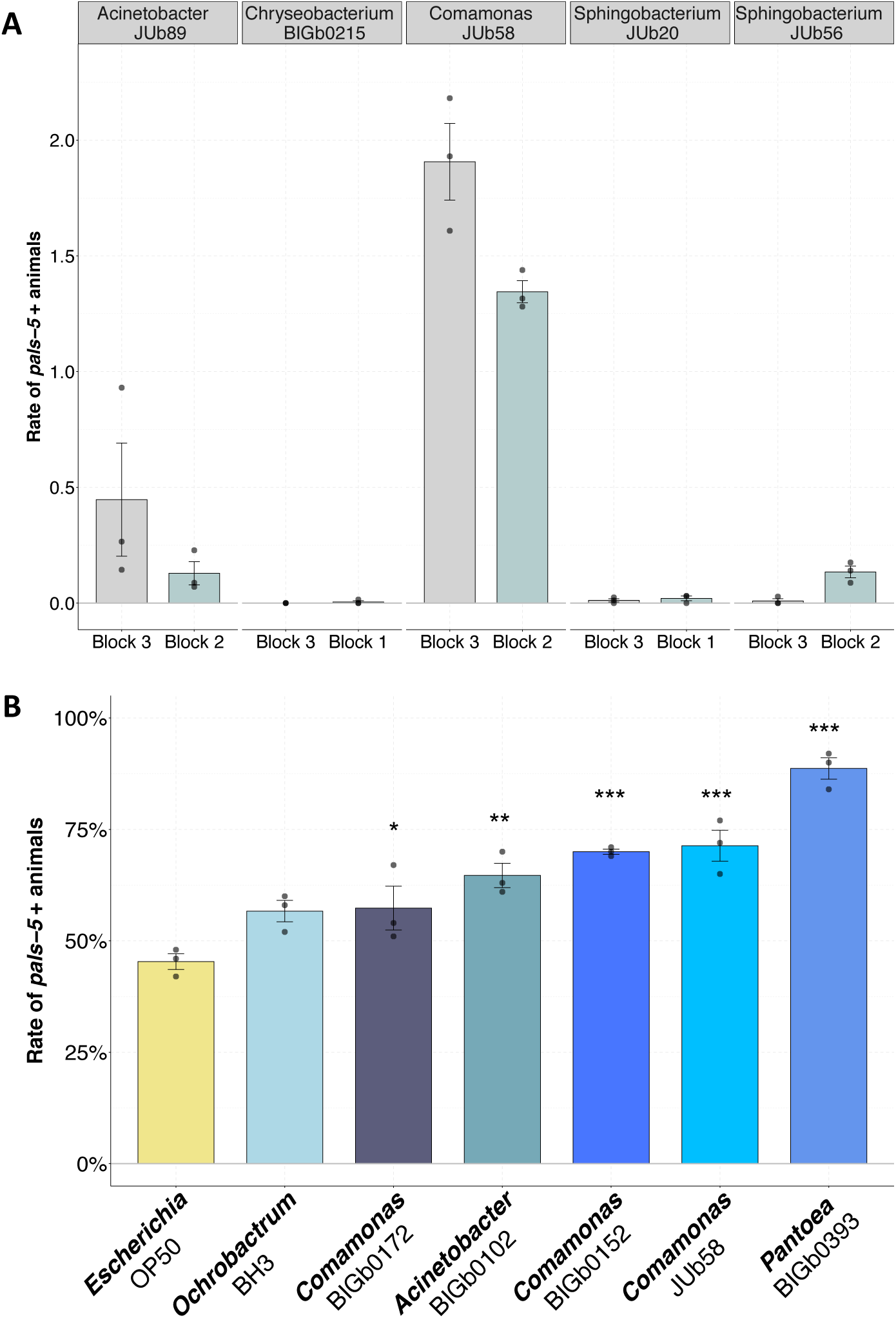
A) The impact of bacterial strains in viral infections is consistent in different experiments. **B)** New experiment with the bacteria that enhanced infection on the initial screening shown in Fig 1. Data are presented as mean ± standard error. Asterisks on the graphs represent values of significance: *** P < 0.001; ** P < 0.01; * 0.01 < P < 0.05; P values higher than 0.05 are not labeled.

**Supplementary Figure 3.**
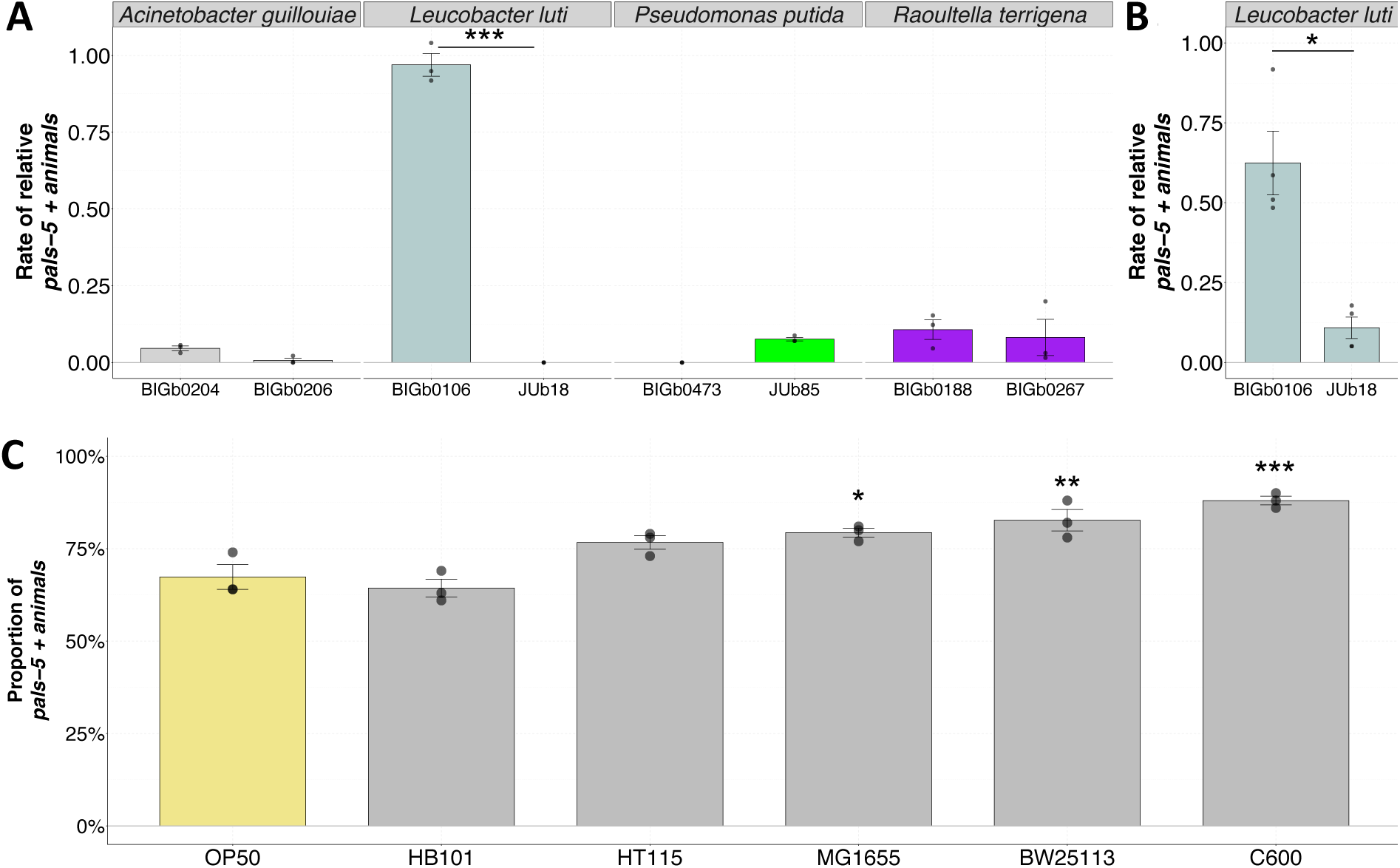
A) Activation of the *pals-5* reporter upon viral infection of animals on different strains of the same natural bacteria species. **B)** Second experiment with *Leucobacter luti*. **C)** Activation of infection reporter upon viral infection of animals on different *E. coli* strains. Data are presented as mean ± standard error. Asterisks on the graphs represent values of significance: *** P < 0.001; ** P < 0.01; * 0.01 < P < 0.05; P values higher than 0.05 are not labeled.

**Supplementary Figure 4.**
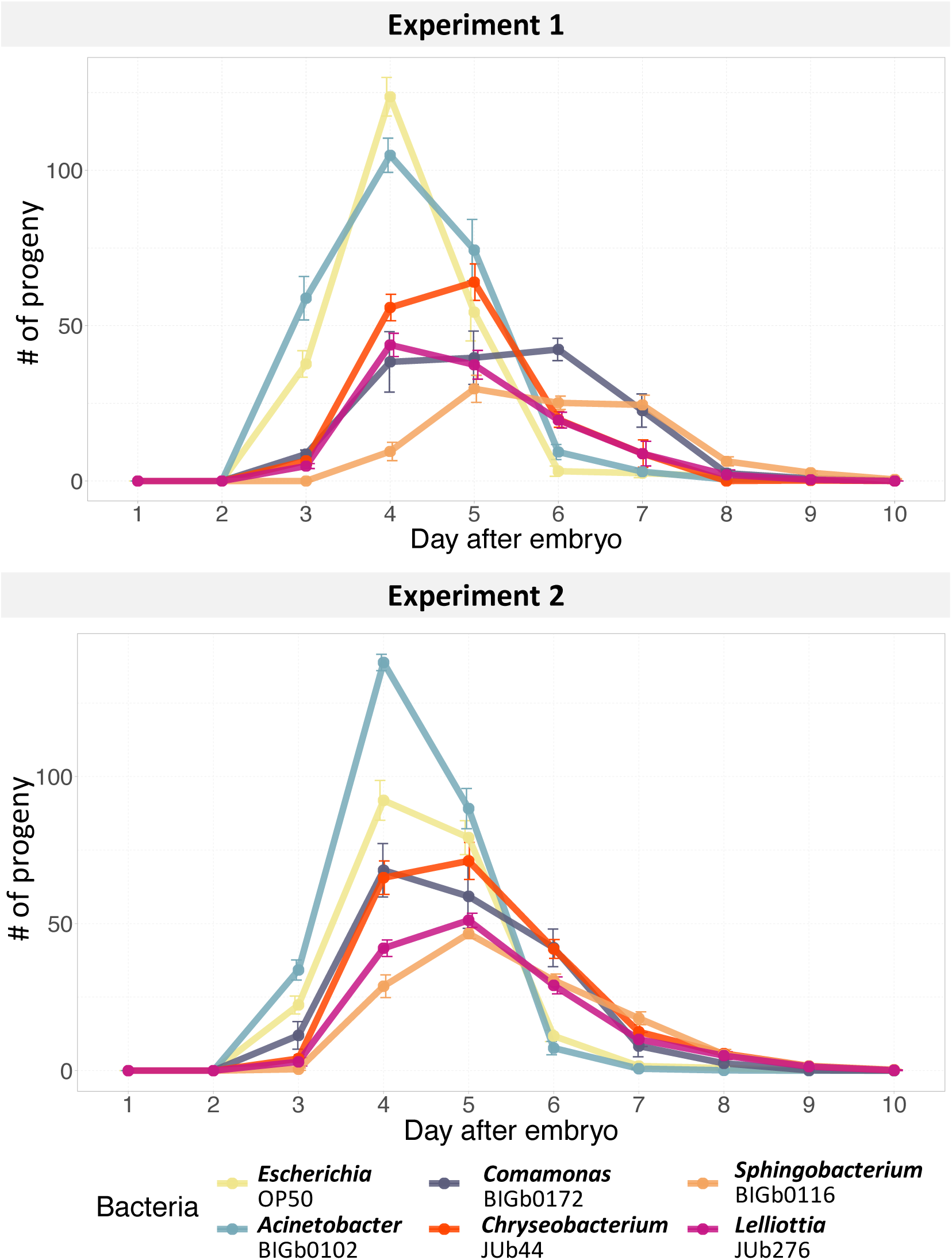
*C. elegans* ERT54 production of progeny per day on selected bacteria, in the absence of virus.

**Supplementary Figure 5.**
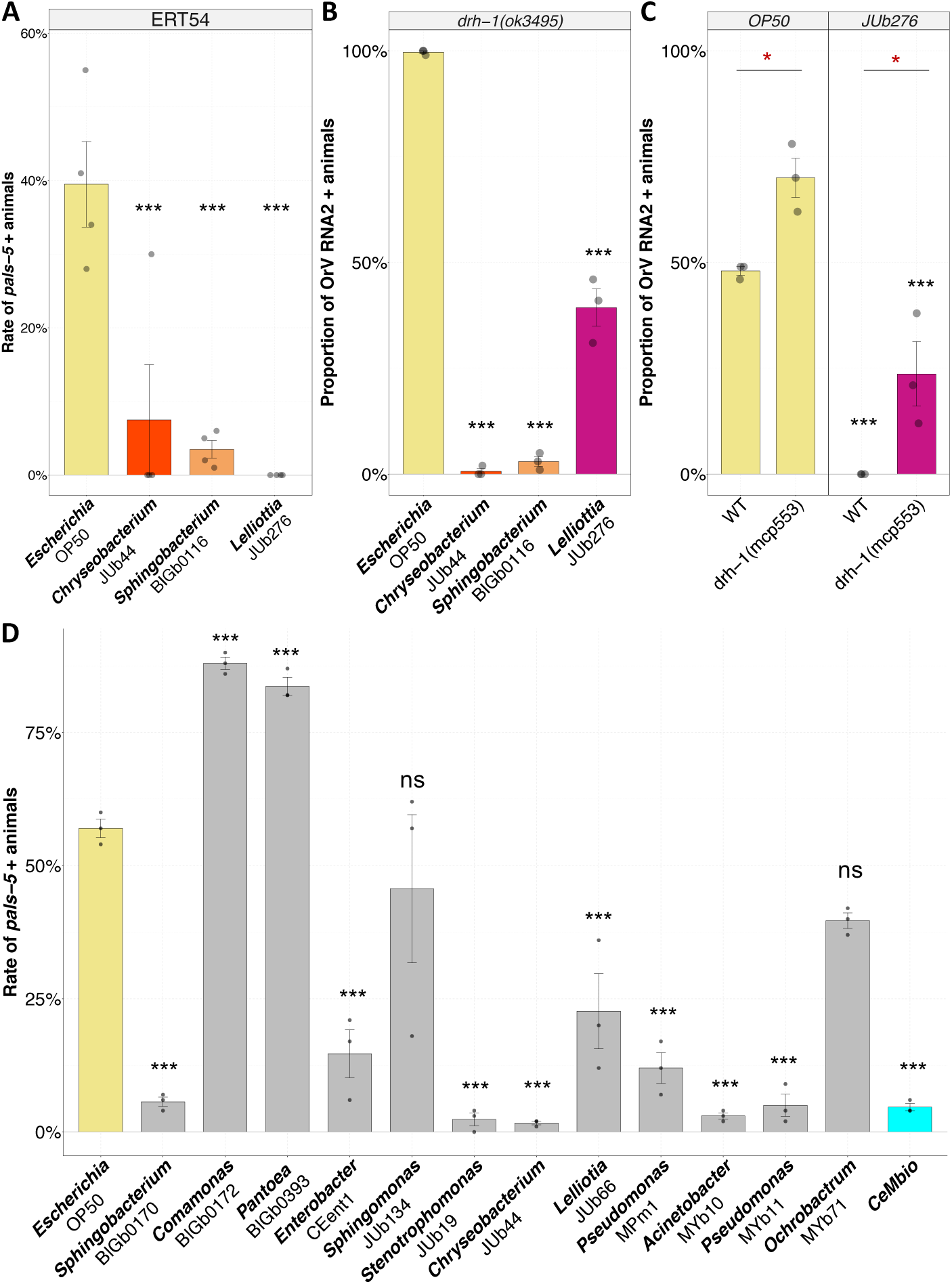
New experiments to confirm results **(A)** Repetition of experiment shown in Figure 3, but chunking transfer after 2 generations. **(B)** Repetition of experiments shown in Figure 6D. **(C)** Confirmation of *drh-1* animals susceptibility to viral infections in JUb276 by testing animals carrying a natural deletion of *drh-1*. **(D)** Activation of the *pals-5* reporter upon viral infection of animals on different single CeMbio strains and the whole CeMbio community. Data are presented as mean ± standard error. Asterisks on the graphs represent values of significance: *** P < 0.001; ** P < 0.01; * 0.01 < P < 0.05; P values higher than 0.05 are labeled as “ns”.

**Supplementary Figure 6.**
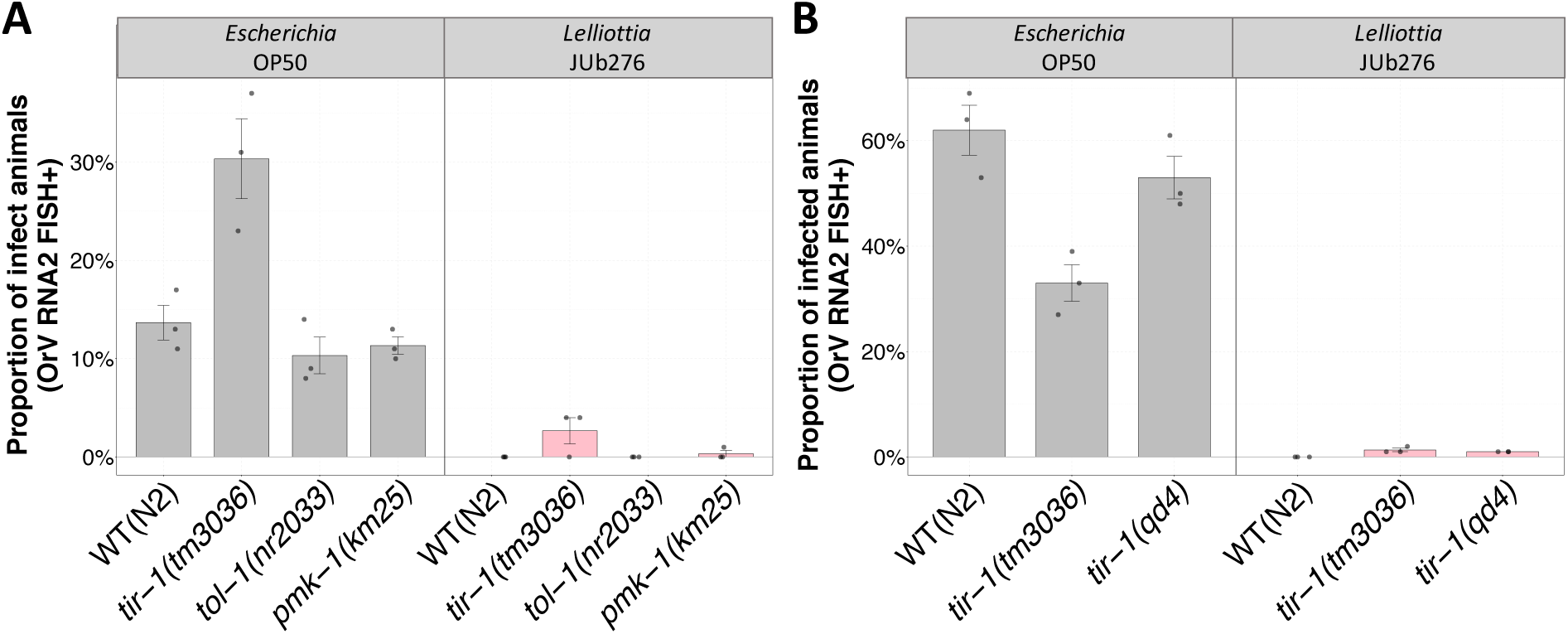
Proportion of viral infected animals on WT or mutants of immune pathways against bacterial infections, on *Escherichia* OP50 or *Lelliottia* JUb276. **(A)** Experiment testing mutants with alterations in different genes involved in the response against bacterial infections. **(B)** Experiment examining *tir-1* mutants. Data are presented as mean ± standard error.

**Supplementary Table 1.** List of bacterial strains used in this work. The *pals-5::GFP* column indicates that *C. elegans* is able to activate the *pals-5::GFP* reporter on all of them after heat shock. For this test, 48 hours-old ERT54 animals were placed at 30 °C for 24 h, being observed right after the 24 h heat shock.

| Strain | Genera | Species | *pals-5::GFP*<br>after heat shock | Reference |
| --- | --- | --- | --- | --- |
| OP50 | *Escherichia* | *Escherichia coli* | Activated | 48 |
| BH3 | *Ochrobactrum* | *Ochrobactrum sp.* | Activated | 61 |
| BIGb0102 | *Acinetobacter* | *Acinetobacter sp.* | Activated | 36 |
| BIGb0106 | *Leucobacter* | *Leucobacter luti* | Activated | 15 |
| BIGb0116 | *Sphingobacterium* | *Sphingobacterium sp.* | Activated | 15 |
| BIGb0138 | *Raoultella* | *Raoultella sp.* | Activated | 15 |
| BIGb0149 | *Raoultella* | *Raoultella sp.* | Not tested | 15 |
| BIGb0152 | *Comamonas* | *Comamonas sp.* | Activated | 15 |
| BIGb0156 | *Scandinavium* | *Scandinavium goeteborgense* | Not tested | 15 |
| BIGb0165 | *Sphingobacterium* | *Sphingobacterium sp.* | Activated | 15 |
| BIGb0170 | *Sphingobacterium* | *Sphingobacterium multivorum* | Activated | 15 |
| BIGb0172 | *Comamonas* | *Comamonas piscis* | Activated | 36 |
| BIGb0188 | *Raoultella* | *Raoultella terrigena* | Not tested | 36 |
| BIGb0204 | *Acinetobacter* | *Acinetobacter sp.* | Not tested | 15 |
| BIGb0206 | *Acinetobacter* | *Acinetobacter guillouiae* | Activated | 15 |
| BIGb0215 | *Chryseobacterium* | *Chryseobacterium nakagawai* | Not tested | 15 |
| BIGb0232 | *Chryseobacterium* | *Chryseobacterium sp.* | Not tested | 15 |
| BIGb0234 | *Serratia* | *Serratia sp.* | Activated | 15 |
| BIGb0236 | *Rahnella* | *Rahnella sp.* | Activated | 15 |
| BIGb0267 | *Raoultella* | *Raoultella terrigena* | Activated | 15 |
| BIGb0273 | *Pseudomonas* | *Pseudomonas brenneri* | Activated | 36 |
| BIGb0359 | *Enterobacter* | *Enterobacter sp.* | Activated | 15 |
| BIGb0383 | *Enterobacter* | *Enterobacter sp.* | Activated | 15 |
| BIGb0393 | *Pantoea* | *Pantoea nemavictus* | Activated | 15 |
| BIGb0399 | *Raoultella* | *Raoultella sp.* | Activated | 15 |
| BIGb0408 | *Pseudomonas* | *Pseudomonas sp.* | Activated | 15 |
| BIGb0435 | *Erwinia* | *Erwinia rhapontici* | Activated | 15 |
| BIGb0473 | *Pseudomonas* | *Pseudomonas putida* | Activated | 15 |
| BIGb0477 | *Pseudomonas* | *Pseudomonas psychrophila* | Not tested | 15 |
| BIGb0494 | *Acinetobacter* | *Acinetobacter johnsonii* | Activated | - |
| BIGb0525 | *Pseudomonas* | *Pseudomonas helmanticensis* | Activated | 15 |
| BIGb0552 | *Buttiauxella* | *Buttiauxella* sp. | Activated | 15 |
| BIGb0611 | *Gluconobacter* | *Gluconobacter cerinus* | Activated | 36 |
| CEent1 | *Enterobacter* | *Enterobacter hormaechei* | Activated | 13 |
| JUb102 | *Providencia* | *Providencia alcalifaciens* | Activated | 15 |
| JUb115 | *Arthrobacter* | *Arthrobacter* sp. | Activated | 15 |
| JUb134 | *Sphingomonas* | *Sphingomonas molluscorum* | Activated | 41 |
| JUb18 | *Leucobacter* | *Leucobacter luti* | Activated | 15 |
| JUb19 | *Stenotrophomonas* | *Stenotrophomonas indicatrix* | Activated | 15 |
| JUb20 | *Sphingobacterium* | *Sphingobacterium* sp. | Activated | 36 |
| JUb202 | *Lelliottia* | *Lelliottia* sp. | Not tested | - |
| JUb21 | *Sphingobacterium* | *Sphingobacterium* sp. | Activated | 15 |
| JUb230 | *Lelliottia* | *Lelliottia* sp. | Activated | - |
| JUb276 | *Lelliottia* | *Lelliottia* sp. | Activated | 37 |
| JUb28 | *Pseudomonas* | *Pseudomonas protegens* | Activated | 15 |
| JUb34 | *Curtobacterium* | *Curtobacterium* sp. | Activated | 15 |
| JUb39 | *Providencia* | *Providencia* sp. | Activated | 15 |
| JUb44 | *Chryseobacterium* | *Chryseobacterium scophthalmum* | Activated | 15 |
| JUb45 | *Rhizobium* | *Neorhizobium* sp. | Activated | 15 |
| JUb52 | *Pseudomonas* | *Pseudomonas* sp. | Activated | 15 |
| JUb53 | *Rahnella* | *Rahnella* sp. | Activated | 15 |
| JUb54 | *Raoultella* | *Raoultella ornithinolytica* | Activated | 15 |
| JUb56 | *Sphingobacterium* | *Sphingobacterium* sp. | Activated | 15 |
| JUb58 | *Comamonas* | *Comamonas* sp. | Activated | 15 |
| JUb65 | *Curtobacterium* | *Curtobacterium flaccumfaciens* | Activated | 36 |
| JUb66 | *Lelliottia* | *Lelliottia amnigena* | Not tested | 15 |
| JUb68 | *Acinetobacter* | *Acinetobacter* sp. | Not tested | 15 |
| JUb7 | *Chryseobacterium* | *Chryseobacterium* sp. | Activated | 15 |
| JUb78 | *Sphingobacterium* | *Sphingobacterium* sp. | Activated | 15 |
| JUb8 | *Delftia* | *Delftia acidovorans* | Activated | 15 |
| JUb85 | *Pseudomonas* | *Pseudomonas putida* | Activated | 15 |
| JUb87 | *Buttiauxella* | *Buttiauxella* sp. | Activated | 15 |
| JUb89 | *Acinetobacter* | *Acinetobacter calcoaceticus* | Activated | 36 |
| JUb96 | *Pseudomonas* | *Pseudomonas* sp. | Activated | 15 |
| MSPm1 | *Pseudomonas* | *Pseudomonas berkeleyensis* | Activated | 41 |
| MYb10 | *Acinetobacter* | *Acinetobacter guillouiae* | Activated | 14 |
| MYb11 | *Pseudomonas* | *Pseudomonas lurida* | Activated | 14 |
| MYb71 | *Ochrobactrum* | *Ochrobactrum vermis* | Activated | 14 |
| CeMBio | Microbial community |  | Activated | 41 |

**Supplementary Table 2.** Nematode strains used in this study.

| Strain | Genotype |
| --- | --- |
| ERT54 | *jyIs8[pals-5p::GFP; myo-2::mCherry]* |
| ERT71 | *jyIs15[F26F2.1p::GFP; myo-2::mCherry]* |
| ERT90 | *zip-1(jy13)* |
| GL302 | *cde-1(rf34)* |
| IG10 | *tol-1(nr2033)* |
| IG685 | *tir-1(tm3036)* |
| JU2508 | *drh-1(ok3495)* |
| JU2624 | *[myo-2::mCherry::unc54; lys3p::eGFP::tbb2]* II in JU1580 |
| KU25 | *pmk-1(km25)* |
| MCP553 | *drh-1(mcp553)* |
| N2 | wild-type reference |
| SX2755 | *sdz-6p::GFP; myo-2::mCherry* |
| WM27 | *rde-1(ne219)* |
| ZD101 | *tir-1(qd4)* |

## REFERENCES

1. Leulier, F., et al. Integrative Physiology: At the Crossroads of Nutrition, Microbiota, Animal Physiology, and Human Health. Cell Metabolism 25, 522–534 (2017).

2. González, R. & Elena, S. F. The Interplay between the Host Microbiome and Pathogenic Viral Infections. mBio 12, e02496–21 (2021).

3. Belkacem, N. et al. *Lactobacillus paracasei* feeding improves immune control of influenza infection in mice. PLoS ONE 12, e0184976 (2017).

4. Fonseca, W. et al. *Lactobacillus johnsonii* supplementation attenuates respiratory viral infection via metabolic reprogramming and immune cell modulation. Mucosal Immunology 10, 1569–1580 (2017).

5. Kim, H. J. et al. Nasal commensal *Staphylococcus epidermidis* enhances interferon-λ- dependent immunity against influenza virus. Microbiome 7, 80 (2019).

6. Stefan, K. L., Kim, M. V., Iwasaki, A. & Kasper, D. L. Commensal Microbiota Modulation of Natural Resistance to Virus Infection. Cell 183, 1312–1324.e10 (2020).

7. Kuss, S. K., et al. Intestinal Microbiota Promote Enteric Virus Replication and Systemic Pathogenesis. Science 334, 249–252 (2011).

8. Robinson, C. M., Jesudhasan, P. R. & Pfeiffer, J. K. Bacterial Lipopolysaccharide Binding Enhances Virion Stability and Promotes Environmental Fitness of an Enteric Virus. Cell Host & Microbe 15, 36–46 (2014).

9. Domínguez-Díaz, C., García-Orozco, A., Riera-Leal, A., Padilla-Arellano, J. R. & Fafutis-Morris, M. Microbiota and Its Role on Viral Evasion: Is It With Us or Against Us? Front Cell Infect Microbiol 9, 256 (2019).

10. Backes, C., Martinez-Martinez, D. & Cabreiro, F. *C. elegans*: A biosensor for host–microbe interactions. Lab Anim 50, 127–135 (2021).

11. Frézal, L. & Félix, M.-A. *C. elegans* outside the Petri dish. eLife 4, e05849 (2015).

12. Schulenburg, H. & Félix, M.-A. The Natural Biotic Environment of *Caenorhabditis elegans*. Genetics 206, 55–86 (2017).

13. Berg, M., et al. Assembly of the *Caenorhabditis elegans* gut microbiota from diverse soil microbial environments. ISME J 10, 1998–2009 (2016).

14. Dirksen, P., et al. The native microbiome of the nematode *Caenorhabditis elegans*: gateway to a new host-microbiome model. BMC Biol 14, 38 (2016).

15. Samuel, B. S., Rowedder, H., Braendle, C., Félix, M.-A. & Ruvkun, G. *Caenorhabditis elegans* responses to bacteria from its natural habitats. Proc Natl Acad Sci USA 113, 3941–3949 (2016).

16. Coolon, J. D., Jones, K. L., Todd, T. C., Carr, B. C. & Herman, M. A. *Caenorhabditis elegans* Genomic Response to Soil Bacteria Predicts Environment-Specific Genetic Effects on Life History Traits. PLoS Genet 5, e1000503 (2009).

17. Cabreiro, F. & Gems, D. Worms need microbes too: microbiota, health and aging in *Caenorhabditis elegans*. EMBO Mol Med 5, 1300–1310 (2013).

18. Watson, E., MacNeil, L. T., Arda, H. E., Zhu, L. J. & Walhout, A. J. M. Integration of Metabolic and Gene Regulatory Networks Modulates the *C. elegans* Dietary Response. Cell 153, 253–266 (2013).

19. Zhang, F., et al. *Caenorhabditis elegans* as a Model for Microbiome Research. Front. Microbiol. 8, 485 (2017).

20. Irazoqui, J. E., et al. Distinct Pathogenesis and Host Responses during Infection of *C. elegans* by P. aeruginosa and S. aureus. PLoS Pathog 6, e1000982 (2010).

21. Kim, Y. & Mylonakis, E. *Caenorhabditis elegans* Immune Conditioning with the Probiotic Bacterium *Lactobacillus acidophilus* Strain NCFM Enhances Gram-Positive Immune Responses. Infect Immun 80, 2500–2508 (2012).

22. Montalvo-Katz, S., Huang, H., Appel, M. D., Berg, M. & Shapira, M. Association with Soil Bacteria Enhances p38-Dependent Infection Resistance in *Caenorhabditis elegans*. Infect Immun 81, 514–520 (2013).

23. King, K. C., et al. Rapid evolution of microbe-mediated protection against pathogens in a worm host. ISME J 10, 1915–1924 (2016).

24. Smolentseva, O., et al. Mechanism of biofilm-mediated stress resistance and lifespan extension in *C. elegans*. Sci Rep 7, 7137 (2017).

25. Kissoyan, K. A. B., et al. Natural *C. elegans* Microbiota Protects against Infection via Production of a Cyclic Lipopeptide of the Viscosin Group. Current Biology 29, 1030–1037.e5 (2019).

26. Félix, M.-A., et al. Natural and Experimental Infection of *Caenorhabditis* Nematodes by Novel Viruses Related to Nodaviruses. PLoS Biol 9, e1000586 (2011).

27. Félix, M.-A. & Wang, D. Natural Viruses of *Caenorhabditis* Nematodes. Annu. Rev. Genet. 53, 313–326 (2019).

28. Tran, T. D. & Luallen, R. J. An organismal understanding of *C. elegans* innate immune responses, from pathogen recognition to multigenerational resistance. Semin Cell Dev Biol S1084952123000605 (2023).

29. González, R. & Félix, M.-A. *Caenorhabditis elegans* immune responses to intracellular pathogens. Dev Comp Immunol **UNDER REVIEW** (2023).

30. Reddy, K. C., et al. An Intracellular Pathogen Response Pathway Promotes Proteostasis in *C. elegans*. Current Biology 27, 3544–3553.e5 (2017).

31. Ashe, A., et al. A deletion polymorphism in the *Caenorhabditis elegans* RIG-I homolog disables viral RNA dicing and antiviral immunity. eLife 2, e00994 (2013).

32. Sowa, J. N., et al. The *Caenorhabditis elegans* RIG-I Homolog DRH-1 Mediates the Intracellular Pathogen Response upon Viral Infection. J Virol 94, e01173–19 (2020).

33. Lažetić, V., et al. The transcription factor ZIP-1 promotes resistance to intracellular infection in *Caenorhabditis elegans*. Nat Commun 13, 17 (2022).

34. Le Pen, J., et al. Terminal uridylyltransferases target RNA viruses as part of the innate immune system. Nat Struct Mol Biol 25, 778–786 (2018).

35. Jiang, H., Chen, K., Sandoval, L. E., Leung, C. & Wang, D. An Evolutionarily Conserved Pathway Essential for Orsay Virus Infection *of Caenorhabditis elegans*. mBio 8, e00940–17 (2017).

36. Zhang, F. et al. Natural genetic variation drives microbiome selection in the *Caenorhabditis elegans* gut. Curr Biol 31, 2603–2618.e9 (2021).

37. Frézal, L., et al. Genome-wide association and environmental suppression of the mortal germline phenotype of wild C. elegans. http://biorxiv.org/lookup/doi/10.1101/2023.05.17.540956 (2023)

38. Bakowski, M. A., et al. Ubiquitin-Mediated Response to Microsporidia and Virus Infection in *C. elegans*. PLoS Pathog 10, e1004200 (2014).

39. Frézal, L., Jung, H., Tahan, S., Wang, D. & Félix, M.-A. Noda-Like RNA Viruses Infecting *Caenorhabditis* Nematodes: Sympatry, Diversity, and Reassortment. J Virol 93, e01170–19 (2019).

40. Grenfell, B. T., Dobson, A. P., & Moffatt, H. K. (Eds.). (1995). Ecology of infectious diseases in natural populations (Vol. 7). Cambridge University Press.

41. Dirksen, P., et al. CeMbio - The *Caenorhabditis elegans* Microbiome Resource. *G3* *Genes|Genomes|Genetics* **10**, 3025–3039 (2020).

42. Ford, S. A. & King, K. C. Harnessing the Power of Defensive Microbes: Evolutionary Implications in Nature and Disease Control. PLoS Pathog 12, e1005465 (2016).

43. Pike, V. L., Lythgoe, K. A. & King, K. C. On the diverse and opposing effects of nutrition on pathogen virulence. Proc. R. Soc. B. 286, 20191220 (2019).

44. Casorla-Perez, L. A., et al. Orsay Virus Infection of *Caenorhabditis elegans* Is Modulated by Zinc and Dependent on Lipids. J Virol 96, e01211–22 (2022).

45. Piqué, N., Berlanga, M. & Miñana-Galbis, D. Health Benefits of Heat-Killed (Tyndallized) Probiotics: An Overview. IJMS 20, 2534 (2019).

46. Zilber-Rosenberg, I. & Rosenberg, E. Role of microorganisms in the evolution of animals and plants: the hologenome theory of evolution. FEMS Microbiol Rev 32, 723–735 (2008).

47. Henry, L. P., Bruijning, M., Forsberg, S. K. G. & Ayroles, J. F. The microbiome extends host evolutionary potential. Nat Commun 12, 5141 (2021).

48. Brenner, S. The genetics of *Caenorhabditis elegans*. Genetics 77, 71–94 (1974).

49. Stiernagle T. Maintenance of *C. elegans*. Wormbook 1,1 (2006).

50. Franz, C. J., et al. Orsay, Santeuil and Le Blanc viruses primarily infect intestinal cells in *Caenorhabditis* nematodes. Virology 448, 255–264 (2014).

51. Katoh, K. & Standley, D. M. MAFFT Multiple Sequence Alignment Software Version 7: Improvements in Performance and Usability. Molecular Biology and Evolution 30, 772–780 (2013).

52. Biegert, A., Mayer, C., Remmert, M., Soding, J. & Lupas, A. N. The MPI Bioinformatics Toolkit for protein sequence analysis. Nucleic Acids Research 34, 335–339 (2006).

53. Trifinopoulos, J., Nguyen, L.-T., von Haeseler, A. & Minh, B. Q. W-IQ-TREE: a fast online phylogenetic tool for maximum likelihood analysis. Nucleic Acids Res 44, 232–235 (2016).

54. Keck, F., Rimet, F., Bouchez, A. & Franc, A. phylosignal: an R package to measure, test, and explore the phylogenetic signal. Ecol Evol 6, 2774–2780 (2016).

55. R Core Team. (2021). R: A language and environment for statistical computing. R Foundation for Statistical Computing. https://www.R-project.org/

56. Hothorn, T., Bretz, F. & Westfall, P. Simultaneous Inference in General Parametric Models. Biom. J. 50, 346–363 (2008).

57. Graves, S., Piepho, H., Selzer, L., Dorai-Raj, S., Potvin, C., & Western, B. (2021). multcompView: Visualizations of Paired Comparisons. R package version 0.1–8. https://CRAN.R-project.org/package=multcompView

58. Bates, D., Mächler, M., Bolker, B., & Walker, S. Fitting linear mixed-effects models using lme4. J Stat Softw 67, 1–48 (2015).

59. Lenth, R. (2020). emmeans: Estimated Marginal Means, aka Least-Squares Means. R package version 1.4.7. https://CRAN.R-project.org/package=emmeans

60. Kassambara, A. (2023). ggpubr: ’ggplot2’ Based Publication Ready Plots. R package version 0.2.4. https://CRAN.R-project.org/package=ggpubr

61. Troemel, E. R., Félix, M.-A., Whiteman, N. K., Barrière, A. & Ausubel, F. M. Microsporidia Are Natural Intracellular Parasites of the Nematode *Caenorhabditis elegans*. PLoS Biol 6, e309 (2008).

